# Deep Recurrent Attention Models for Histopathological Image Analysis

**DOI:** 10.1101/438341

**Authors:** Alexandre Momeni, Marc Thibault, Olivier Gevaert

## Abstract

Histopathology defines the gold standard in oncology. Automatic analysis of pathology images could thus have a significant impact on diagnoses, prognoses and treatment decisions for cancer patients. Recently, convolutional neural networks (CNNs) have shown strong performance in computational histopathology tasks. However, given it remains intractable to process pathology slides in their entirety, CNNs have traditionally performed inference on small individual patches extracted from the image. This often requires a significant amount of computation and can result in ignoring potentially relevant spatial and contextual information. Being able to process larger input patches and locating discriminatory regions more efficiently could help improve both computational and task specific performance. Inspired by the recent success of Deep Recurrent Attention Models (DRAMs) in image recognition tasks, we propose a novel attention-based architecture for classification in histopathology. Similar to CNNs, DRAMs have a degree of translation invariance built-in, but the amount of computation performed can be controlled independently from the input image size. The model is a deep recurrent neural network trained with reinforcement learning to attend to the most relevant areas of large input patches. We evaluate our model on histological and molecular subtype classification tasks for the glioma cohorts of The Cancer Genome Atlas (TCGA). Our results suggest that the DRAM has comparable performance to state-of-the-art CNNs despite only processing a select number of patches.

## I. INTRODUCTION

Analysis of histopathology images is a critical step in oncology as interpretation of these images by pathologists is the gold standard for diagnosis, prognosis and treatment. Interpretation of histopathology images largely consists of careful microscopic examination of hematoxylin and eosin (H&E) stained tissue sections. This can be a tedious, time-consuming and sometimes subjective task. Advances in slide scanning technology and reductions in cost of digital storage capacity have enabled the widespread adoption of digital pathology over the past decade [1]. More recently, the dramatic increase in computational power and breakthroughs in machine learning have fueled research into automated analysis of histopathology images [2]. These developments have together led to the rapid emergence of computational histopathology. Most recent works have successfully leveraged Convolutional Neural Networks (CNNs) for tasks such as detection, image segmentation and diagnosis, highlighting the effectiveness and relevance of learned features in complex images such as histopathology slides [3], [4], [5].

It is important to highlight however that several challenges have to be overcome before deep learning can be applied succesfully in the field of digital pathology. In addition to the scarcity of training samples typical for medical image datasets, the key challenges that hinder a deep learning approach include the lack of ground-truth localized annotations and the extremely large size of digital pathology slides (often in the order of gigapixels). The former challenge implies histopathology image analysis (HIA) belongs to the field of Weakly Supervised Learning (WSL). The latter challenge implies it is computationally intractable to process a slide in its entirety. Thus, any modeling approach needs to be time and memory efficient whilst extracting as much information as possible out of the image. The popular approach adopted to deal with this has been to formulate HIA as a Multiple Instance Learning (MIL) problem: instead of treating images as a single instance, we instead assume it represents a bag of instances [6], [27], [28]. This consists in dividing the histopathology slides into small high resolution patches, sampling randomly from these patches and applying patch level CNNs. The MIL framework is then used to combine patch level predictions intelligently and make an overall slide prediction.

Despite their general success in HIA, these methodologies have several drawbacks. Firstly, given each slide may contain a few thousand patches, a non-negligible computational burden may be associated to processing a representative proportion of the entire slide. Furthermore, given there is inherent loss of contextual and spatial information when dealing with randomly sampled patches, the final aggregation step can be particularly difficult to generalize and may be suboptimal.

In a study on pathologist eye movement Brunye et al. discussed findings on expert visual diagnostic procedures [7]. Indeed, they highlighted that visual attention to salient but diagnostically irrelevant image features may underlie the inherently different viewing behavior of novices versus experts while interpreting digitized breast pathology slides. Thus, attentional biases in highly trained professionals may be a core skill to quickly identify relevant areas in slides by focusing on a few select regions.

We propose to mimic the pathologists process by using Deep Recurrent Attention Models (DRAMs), initially proposed by Mnih et al. [8]. DRAMs are inspired by the way humans perform visual recognition tasks. These networks dynamically attend to the most informative areas of an image, only processing these regions. The architecture is independent of input image size, allowing for much larger patch size and limiting computational cost. In this paper, we propose a model based on DRAMs to distinguish glioma subtypes using data from The Cancer Genome Atlas (TCGA).

Gliomas are the most common type of adult brain tumors with variable response to therapy, clinical course, and prognoses. In 2016, the World Health Organization (WHO) released its new Classification of Tumors of the Central Nervous System [9], this integrated new genetic and molecular information to explain tumor heterogeneities. We evaluated our algorithm on the following tasks: tumor grade (i.e. lower grade vs. glioblastoma), IDH1 mutation status, 1p/19q codeletion, and MGMT methylation status. Thus we show that it is possible to gain rapid and accurate classification of these crucial variables from a pathology image which could assist clinicians into better and more timely decision making.

## II. RELATED WORK

### A. Computational Histopathology

Convolutional Neural Networks (CNNs) were first used in HIA for mitosis detection in Whole Slide Image (WSI) [3]. This work was instrumental in demonstrating the effectiveness and relevance of learned features in complex images such as histopathology slides. The application of CNNs has since become widespread in histopathology tasks. Many works make use of a multiple instance learning (MIL) framework [6] to aggregate patch level inference into a global image level classification. Their contribution often lies in presenting novel aggregation methodologies to improve the transition from local to global inference. For example, Hou et al. [5] proposed an EM-based method to identify discriminative patches in high resolution images automatically, whilst Ilse et al. [23] recently presented an attention-based framework assigning larger weights to key instances. There have also been focused efforts on using latent spaces to improve inference. Zanjani et al. [21] successfully applied Conditional Random Fields (CRFs) over latent spaces of a trained CNN in order to jointly assign labels to the patches. This led to performing inference on a wider context rather than appearance of individual patches. Our work goes beyond CNN architectures and integrates Visual Attention techniques. To our knowledge, there have been no previous attempts to applying such *hard attention* architectures to computational histopathology problems.

### B. Visual Attention Models

Initially motivated by the need to reduce the computational cost associated with CNNs, attention-based models have demonstrated state-of-the-art performance on a variety of image recognition tasks in recent years [8], [15], [16]. The original RAM proposed by Mnih et al. takes inspiration from the way humans perform visual recognition [8]. It uses a recurrent neural network to sequentially integrate information extracted from *glimpses* cropped out of an image. Given the cropping operation is non differentiable, the authors resort to reinforcement learning to train the model via the REINFORCE rule [17]. RAM significantly outperformed CNN architectures with a comparable number of parameters on several datasets including the MNIST dataset [18], highlighting the potential of this architecture. Ba et al. extended this line of work with the Deep Recurrent Visual Attention Model (DRAM) [15], achieving better performance than RAM on MNIST classification and producing state-of-the-art results on SVHN recognition [19]. More recently, Jaderberg et al. [20] proposed Spatial Transformer Networks (STN), a trainable module very similar in nature to a recurrent attention model, which has also shown state-of-the-art results on the same tasks as DRAM.

Our model is most similar to the architectures described by Mnih et al. [8] and Ba et al. [15].

## III. METHODS

In this section we present the data, our pre-processing pipeline and provide a description of the model. We also discuss model evaluation.

### A. Data

TCGA [32] provides a rich dataset of H&E stained tissue Whole Slide Image (WSI) for a variety of cancer types. For our experiments, we focus on patients from the glioma cohorts, namely TCGA-LGG (Lower Grade Glioma) and TCGA-GBM (Glioblastoma). We download and preprocess 710 glioma cases (339 LGG & 371 GBM patients respectively) from the GDC portal. Along with the slides, we used labels provided in the Supplementary Materials of [25] for our classification tasks.

### B. Whole Slide Image Pre-processing

We perform the pre-processing steps on the highest slide resolution available (40x magnification).

#### Region of Interest

Tissue segmentation is necessary given there are large areas of white background space in histopathology images which are irrelevant for analysis. We follow a threshold based segmentation method to automatically detect the foreground region. In particular, we first transform the image from RGB to HSV color space and apply Otsu’s method [12] to find the optimal threshold in each channel. The masks are then combined to compute the final tissue segmentation.

#### Tiling

We tile the tissue region extracted from the original slides into large patches. For our experiments we use tiles of 4096 × 4096 pixels, which is the largest tile size we can feasibly perform computation on. The median patient in our cohort has 6 large tiles.

#### Color Normalization

Stain normalization is essential given the results from the staining procedure can vary greatly. Moreover, differences in slide scanners or staining protocol can materially impact stain color, which in turn can affect algorithm performance. Many methods have been introduced to overcome this well defined problem, including sophisticated end-to-end deep learning solutions [13]. For simplicity, we resort to a histogram equalization algorithm as proposed in [10].

Preprocessing pathology slides is a computationally expensive and time consuming task. In their work on histopathology, Ruiz et al. [26] used a GPU to reduce the execution time. We follow their methodology to perform our preprocessing tasks in under 3 hours using multi-threading and program optimization.

### C. Deep Recurrent Attention Model

A Recurrent Attention Model (RAM) processes information in a sequential manner, building up a dynamic internal representation of an environment, which in this paper is the histopathology tile. At each time step *t*, the model focuses selectively on a given location in the large patch called a *glimpse*, then the model extracts features from this glimpse, updates its internal state and chooses the next location to attend. This process is repeated for a fixed number of steps, during which the model incrementally combines the information in a coherent manner. The general architecture can be broken down into a number of sub-components that consist of a multi-layer neural network, where each sub-component maps some input vector into an output vector (Fig. 2). The next section provides a detailed description of each of these sub-components.

**Fig. 1.**
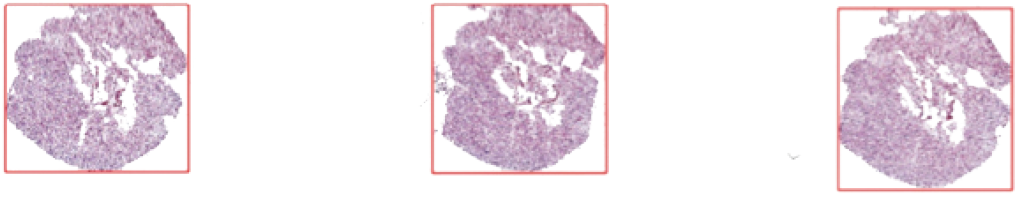
Foreground detection result illustrated by red bounding box. An Otsu method is applied to find the optimal threshold for each channel after the image is converted from RGB to HSV.

**Fig. 2.**
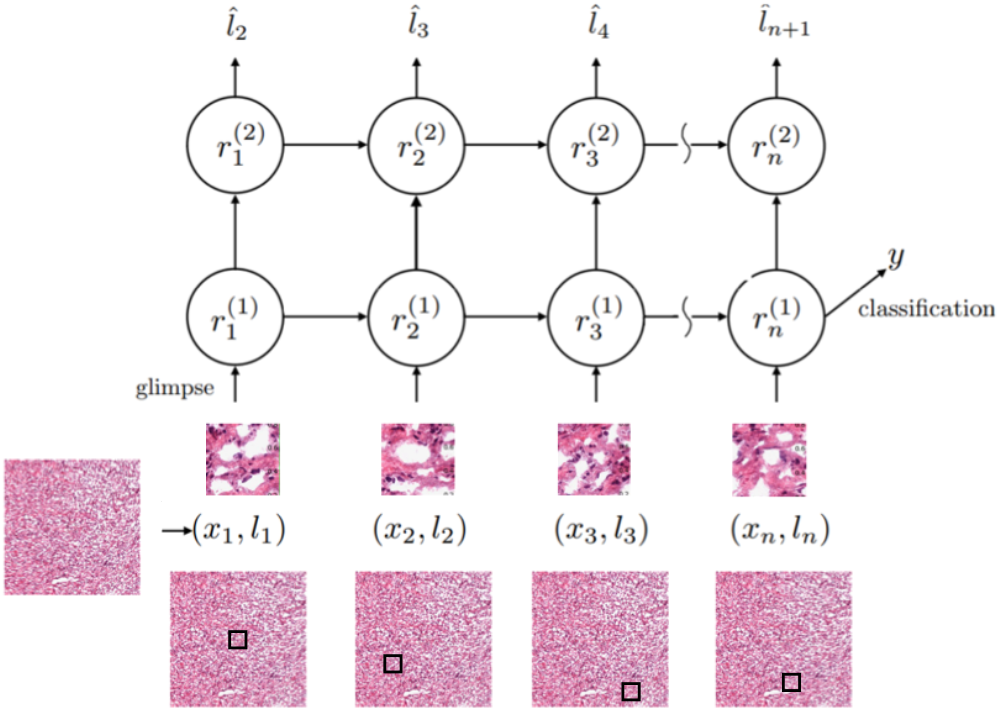
The Deep Recurrent Attention Model. The input is a tile from the histopathology image. At each time step, the model focuses selectively on a given location, extracts features, updates its internal state and chooses the next location to attend. The process is repeated for a fixed number of steps, during which the model incrementally combines the information in a coherent manner to produce a final classification.

**Fig. 3.**
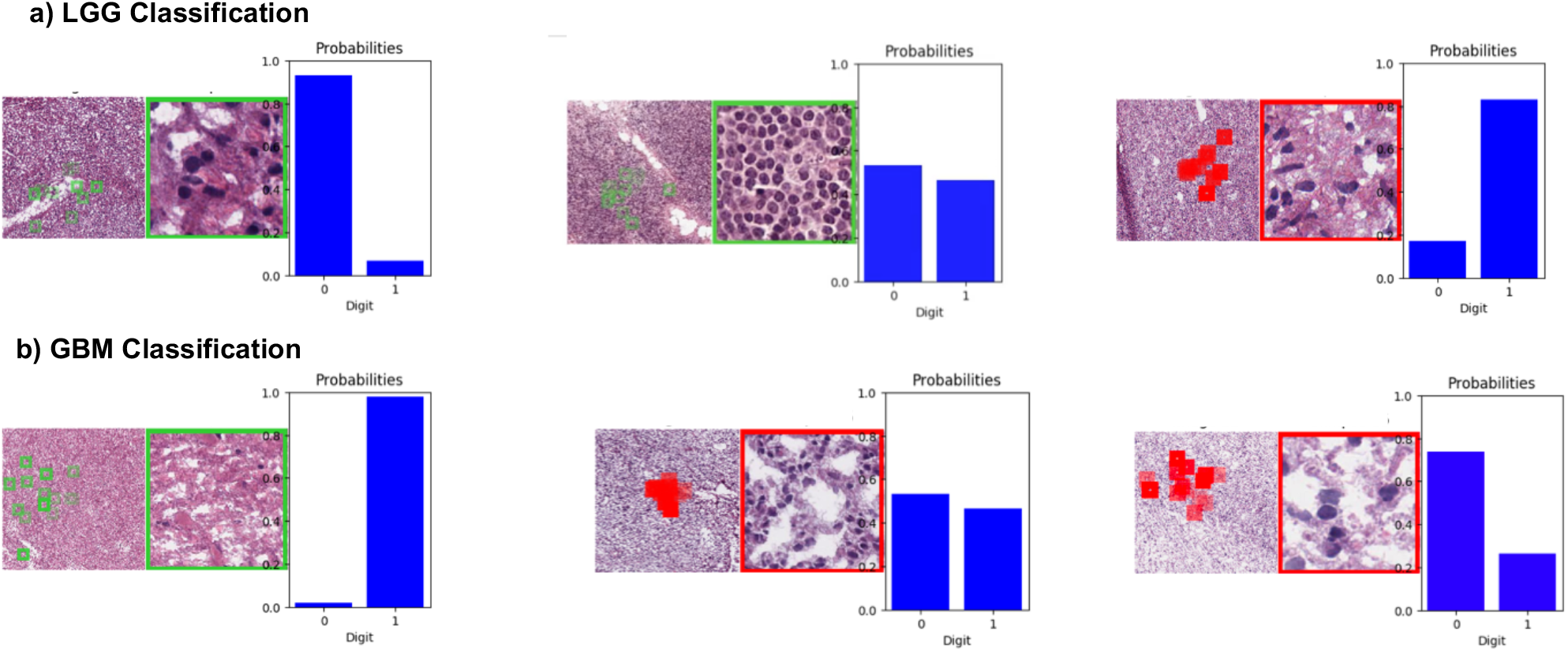
An illustration of the DRAM glimpse map and classification confidence after 16 glimpses.

#### Glimpse Network

At each time step *t*, the glimpse network receives as inputs a location tuple *l_t_* and the corresponding extracted glimpse *x_t_*. The glimpse *x_t_* is sent through three convolutional layers, max pooling, and a fully connected layer to encode a *what* feature vector 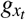. Separately, the location tuple *l_t_* is sent through a fully connected layer to encode a *where* feature vector 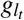. Next, the *what* and *where* feature vectors are multiplied element-wise as initially proposed in [14] and passed through a rectified linear unit activation function to get the final output feature vector *g_t_*.

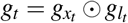

In all our experiments, we initialize *l*_1_ with (0, 0), i.e. the glimpse at the center of the image.

#### Core Network

The core of the model is a two layer Recurrent Neural Network that contains stacked LSTM units [24]. At each time step *t*, the network receives as input *g_t_*, the glimpse network’s feature vector, and encodes two features vectors 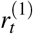 and 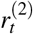:

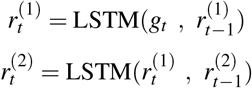

We use LSTM units because of their ability to learn longrange dependencies and stable learning dynamics.

#### Location Network

Using a fully connected layer that takes input feature vector 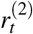 from the top recurrent layer, the location network computes the next glimpse location *l_t_*_+1_. Ideally, the model should learn to look at locations that are relevant for classification. However, the Location Network is non-differentiable, which means standard back-propogation techniques are not applicable. As proposed in [8] [15], we use a policy gradient method in the form of REINFORCE algorithm [17] to train this part of the model.

#### Classification Network

The classification network takes the final feature vector 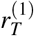 from the lower recurrent layer and outputs a prediction.

As proposed by Mnih et al. [8], we used a hybrid supervised loss in training to optimize the network. In particular, classification loss was defined as cross-entropy between the final prediction and the ground-truth label; REINFORCE [17] was used to train the location network.

### D. Model building and evaluation

For all experiments, we use stratified and shuffled splits to separate the data into training (70%), validation (10%) and testing (20%) sets, using early stopping to avoid overfitting. We also randomly rotated tiles by multiples of 90 degrees and flipped them vertically and horizontally. During training we sample balanced batches, meaning batches contained the same number of patients from each class. We set the glimpse size to 256 × 256 pixels, given this is a common patch size in CNN architectures. We assigned the slide level label to every tile and perform tile-level classification to encourage the network to attend discriminatory regions. However, during testing, we used the average class probabilities over all tiles to classify a slide. We use the following metrics to evaluate all models: classification accuracy, precision, recall, F-score, and area under the receiver operating characteristic curve (AUC).

## IV. RESULTS

We select glioma tumor grade as an initial classification task to solve in order to benchmark our model vs. existing methods. In particular, we consider the results reported by Barker et al. [31], Rubin et al. [30], and Hou et al. [5] for LGG vs. GBM classification on the TCGA dataset. We then apply the method to molecular data.

### A. Prediction of tumor grade

For histological subtype classification (Table I), in order to compare the performance to state-of-the-art results convolutional networks, we use classification accuracy as our performance metric. Our method achieves results statistically equivalent to state-of-the-art reported results for LGG vs. GBM classification (DRAM 24 glimpses, Table I). The model with 24 glimpses has an accuracy, precision, recall, F-Score and AUC of 94% (90%-98% confidence interval), 95%, 93%, 90% and 93% respectively.

**TABLE I.** CLASSIFICATION OF GLIOMA GRADE WITH DRAM AND COMPARISON WITH STATE OF THE ART METHODS

| METHOD | ACCURACY |
| --- | --- |
| Barker et al. | 93% |
| Rubin et al. | 96% |
| Hou et al. | 97% |
| DRAM (4 glimpses) | 77% |
| DRAM (8 glimpses) | 83% |
| DRAM (16 glimpses) | 92% |
| DRAM (24 glimpses) | 94% |

Furthermore, we notice that in every misclassified GBM case at least one tile had been correctly classified. Thus, in cases where the class distribution is highly unbalanced it may be appropriate to use a more sophisticated aggregation strategy.

### B. Prediction of molecular characteristics

For molecular subtype classification, our results suggest it is feasible to use the DRAM approach to classify clinically relevant molecular alterations in both low and high-grade gliomas (Table II). In particular, IDH mutation, MGMT methylation and 1p/19q codeletion are classified correctly with accuracies of 79%, 77% and 71% respectively as well as AUCs of 86%, 75% and 76%. These preliminary results suggest a link with the tissue image which could be explored in further work.

**TABLE II.**
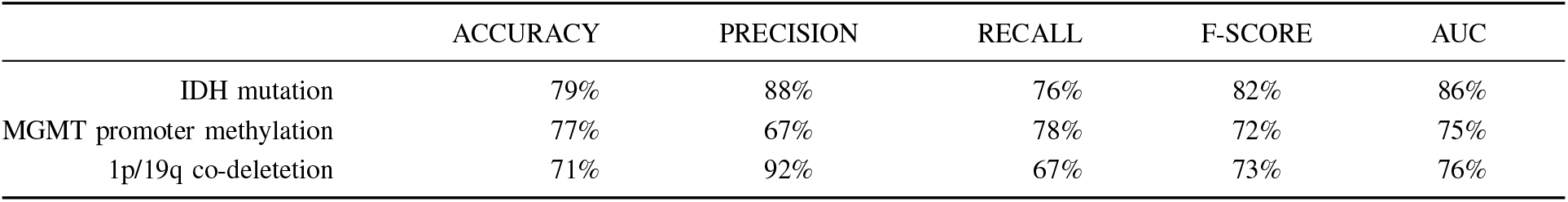
CLASSIFICATION ACCURACY FOR DRAM PREDICTING MOLECULAR CHARACTERISTICS OF GLIOMA

|  | ACCURACY | PRECISION | RECALL | F-SCORE | AUC |
| --- | --- | --- | --- | --- | --- |
| IDH mutation | 79% | 88% | 76% | 82% | 86% |
| MGMT promoter methylation | 77% | 67% | 78% | 72% | 75% |
| 1p/19q co-deletion | 71% | 92% | 67% | 73% | 76% |

## V. DISCUSSION

We have shown that DRAMs can be useful and informative for classification tasks in histopathology. Our results suggest that by selectively choosing where to focus, DRAMs can achieve a performance comparable to state-of-the-art algorithms whilst limiting the computation involved in histopathological image analysis. Our method takes just over two seconds on average to process an entire 4096 × 4096 tile, which is 5 to 10 times larger than the standard tile size. Furthemore, the aggregated classification is performed on 144 glimpses vs. hundreds or thousands of patches in the classical CNN histopathology classifier such as in [5]. Finally, we notice the usefulness of this method in locating clinically significant regions, all while providing a confidence level, making the analysis more interpretable.

However, in future work we would like to explore a number of improvements that can be made to the model. First, our model can be augmented by adding a negative reward for each glimpse it takes, forcing the network to decide when to stop taking glimpses. Second, we could extend the model’s spatial attention mechanism to incorporate multiple resolutions to mimic a pathologist’s process even more accurately.

Finally, in certain slides, the class distribution may be highly unbalanced in favor of the less clinically significant outcome, which suggests more sophisticated aggregation strategies may be beneficial. Incorporating these changes may improve the robustness as well as the speed of convergence of the model.

## REFERENCES

[1] Madabhushi, A. & Lee, G. Image analysis and machine learning in digital pathology: challenges and opportunities. Med. Image Anal. 33, 170175 (2016).

[2] Yann LeCunn, Yoshua Bengio, and Geoffrey Hinton. Deep learning. Nature, 521(7553):436444, 2015.

[3] Dan C. Ciresan, Alessandro Giusti, Luca M. Gambardella, and Jurgen Schmidhuber. Mitosis detection in breast cancer histology images with deep neural networks. In Int. Conf. on Medical Image Computing and Computer-Assisted Intervention, pp. 411418, 2013.

[4] Yun Liu and Krishna Kumar Gadepalli and Mohammad Norouzi and George Dahl and Timo Kohlberger and Subhashini Venugopalan and Aleksey S Boyko and Aleksei Timofeev and Philip Q Nelson and Greg Corrado and Jason Hipp and Lily Peng and Martin Stumpe, Detecting Cancer Metastases on Gigapixel Pathology Images, arXiv preprint arXiv:1703.02442, 2017.

[5] Le Hou, Dimitris Samaras, Tahsin M. Kurc, Yi Gao, James E. Davis, and Joel H. Saltz, Patch-based Convolutional Neural Network for Whole Slide Tissue Image Classification, In Proc. IEEE Conf. on Computer Vision and Pattern Recognition, pp. 24242433, June 2016.

[6] Thomas G. Dietterich, Richard H. Lathrop, and Tomas Lozano-Perez. Solving the multiple instance problem with axis-parallel rectangles. Artificial Intelligence, 89(12):3171, 1997.

[7] Tad T. Brunye, Patricia A. Carney, Kimberly H. Allison, Linda G. Shapiro, Donald L. Weaver, Joann G. Elmore, Eye Movements as an Index of Pathologist Visual Expertise: A Pilot Study, PloS one, 9(8):e103477, 2014

[8] Mnih, Volodymyr, Heess, Nicolas, Graves, Alex, and Kavukcuoglu, Koray. Recurrent models of visual attention. arXiv preprint arXiv:1406.6247, t2014.

[9] Louis DN, Perry A, Reifenberger G, et al. The 2016 World Health Organization Classification of Tumors of the Central Nervous System: a summary. Acta Neuropathol 2016;131:80320 doi:10.1007/s00401-016-1545-1 pmid:27157931

[10] Dennis Nikitenko, Michael A. Wirth, and Kataline Trudel. Applicability of white-balancing algorithms to restoring faded colour slides: An empirical evaluation. Journal of Multimedia, 3(5):918, 2008.

[11] Ciompi, F., Geessink, O., Bejnordi, B.E., de Souza, G.S., Baidoshvili, A., Litjens, G., van Ginneken, B., Nagtegaal, I., van der Laak, J.: The importance of stain normalization in colorectal tissue classification with convolutional networks. arXiv preprint arXiv:1702.05931 (2017).

[12] N. Otsu. A threshold selection method from gray-level histograms. IEEE Transactions on Systems, Man, and Cybernetics, 9(1):6266, 1979

[13] StainGAN: Stain Style Transfer for Digital Histological ImagesvMT Shaban, C Baur, N Navab, S Albarqouni arXiv preprint arXiv:1804.01601

[14] Hugo Larochelle and Geoffrey E. Hinton. Learning to combine foveal glimpses with a third-order boltzmann machine. In NIPS, 2010.

[15] J. Ba, V. Mnih, and K. Kavukcuoglu. Multiple object recognition with visual attention. arXiv preprint arXiv:1412.7755, 2014.

[16] K. Xu, J. Ba, R. Kiros, A. Courville, R. Salakhutdinov, R. Zemel, and Y. Bengio. Show, attend and tell: Neural image caption generation with visual attention. arXiv preprint arXiv:1502.03044, 2015.

[17] R.J. Williams. Simple statistical gradient-following algorithms for connectionist reinforcement learning. Machine Learning, 8(3):229256, 1992.

[18] yann.lecun.com/exdb/mnist/

[19] ufldl.stanford.edu/housenumbers/

[20] Jaderberg M., Simoyan K., Zisserman A., et al., Spatial transformer networks. In Advances in Neural Information Processing Systems (2015), pp. 2017202

[21] Farhad Ghazvinian Zanjani; Svitlana Zinger; Peter H. N. de With, Cancer detection in histopathology whole-slide images using conditional random fields on deep embedded spaces, Proceedings Volume 10581, Medical Imaging 2018: Digital Pathology; 105810I (2018)

[22] Pierre Courtiol, Eric W. Tramel, Marc Sanselme, Gilles Wainrib, Classification and Disease Localization in Histopathology Using Only Global Labels: A Weakly-Supervised Approach, arXiv preprint arXiv:1802.02212

[23] Maximilian Ilse, Jakub M. Tomczak, Max Welling, Attention-based Deep Multiple Instance Learning, arXiv preprint arXiv:1802.04712

[24] Hochreiter, Sepp and Schmidhuber, Jurgen. Long short-term memory. Neural computation, 9(8):17351780, 1997.

[25] Molecular Profiling Reveals Biologically Discrete Subsets and Pathways of Progression in Diffuse Glioma

[26] Ruiz et al., 2007 A. Ruiz, O. Sertel, M. Ujaldon, U. Catalyurek, J. Saltz, M. Gurcan Pathological image analysis using the GPU: stroma classification for neuroblastoma Proceedings of IEEE International Conference on Bioinformatics and Biomedicine, BIBM 2007. (2007), pp. 7888

[27] Z. Jia, X. Huang, E.I.-C. Chang, Y. Xu, Constrained deep weak supervision for histopathology image segmentation, arXiv preprint, arXiv:170100794, 2017

[28] Y. Xu, T. Mo, Q. Feng, P. Zhong, M. Lai, and E. I.-C. Chang, Deep learning of feature representation with multiple instance learning for medical image analysis, in ICASSP, 2014, pp. 16261630

[29] Nesterov, Y. A method of solving a convex programming problem with convergence rate O(1/sqr(k)). Soviet Mathematics Doklady, 27:372376, 1983.

[30] Mehmet Gunhan Ertosun, Daniel L. Rubin, Automated Grading of Gliomas using Deep Learning in Digital Pathology Images: A modular approach with ensemble of convolutional neural networks, AMIA Annu Symp Proc. 2015; 2015: 18991908. 2015.

[31] Automated classification of brain tumor type in whole-slide digital pathology images using local representative tiles, Barker, Jocelyn. et al. Medical Image Analysis, Volume 30, 60-71

[32] portal.gdc.cancer.gov

